# Communication Breakdown and Evolution of the Cancer Cell

**DOI:** 10.1101/2025.08.15.670478

**Authors:** Nemati Fard Lorenzo Amir, Arora Chakit, Miglionico Pasquale, Varisco Martina, Bisceglia Luisa, Vukotic Ranka, Raimondi Francesco

## Abstract

We studied cell-cell interactions (CCIs) in large-scale transcriptomic datasets, which showed higher co-expression in cancer compared to healthy tissues. CCIs are more co-expressed than any other type of intracellular interaction and, likewise, they are the protein-protein interaction (PPI) class that is most co-evolved in sequenced genomes. Similar trends of stricter regulation and evolutionary pressure are observed when comparing extracellular versus intracellular interactions mediated by G protein Coupled Receptors (GPCRs), whose ligand interactions are also characterized by a higher mutational burden in later tumor stages when considering somatic mutations associated with tumor clonal evolution. CCIs undergo the most extensive rewiring of their tumor co-expression networks relative to healthy tissues, more so than any other PPI type, with a set of CCI hubs highly conserved across multiple tumor tissues, and a higher diversity on healthy ones. Cancer rewiring is also associated with the formation of recurrent circuits of co-expressed CCI pairs, represented by enriched network motifs such as triad or tetrad cliques. These act as integrative hotspots to facilitate the crosstalk of distinct processes and the interaction of the cancer cell with its tumor microenvironment (TME). Remarkably, many CCI circuits are significantly associated with patient survival and are predictive of patient response to immunotherapy. CCI circuits mapping to allograft rejection and inflammatory response inform immunotherapy response prediction, while those related to epithelial-mesenchymal transition are associated with poorer prognosis. Overall, we show that CCIs expression signatures could be effectively exploited to stratify patients and, at the same time, they highlight new combination therapeutic opportunities in personalized medicine settings.

## 2 Introduction

Cell-cell interactions (CCIs) are essential components that orchestrate the life of a cell in the social network of a multicellular organism (Su et al., 2024; Armingol et al., 2021). The functions entailed by CCIs represented major work horses for the evolution of multicellular organisms (Li et al., 2025; Combarnous and Nguyen, 2020; Du et al., 2015; De Mendoza et al., 2014), given their essential roles in cellular growth, development, differentiation as well as tissue and organ formation, maintenance, and physiological regulation (Su et al., 2024). CCIs mediate the interactions of membrane receptors with diverse molecules, ranging from ions, metabolites and hormones to secreted peptides, proteins, integrins, junction and structural proteins. Some molecules, such as adhesion proteins, have structural or scaffolding roles, whereas secreted ligands such as hormones, neurotransmitters, growth factors, chemokines or cytokines mediate cell-cell communication. Receiver cells integrate a multitude of such incoming CCIs through membrane receptors and trigger downstream signaling and transcriptional program regulation, ultimately responding to the microenvironment (Su et al., 2024; Armingol et al., 2021). The cell-cell communication network is aberrantly regulated in the tumor microenvironment (TME). Tumor cells foster their growth and proliferation by interacting with surrounding normal cells, immune cells, fibroblasts and endothelial cells within the TME. Cancer cells engage with TME through extensive chemical and physical interactions that are instrumental for multiple tumor characteristics, such as dysregulated extracellular matrix (ECM), sustainment of proliferative signals, inhibition of apoptosis, invasion and metastasis, metabolic dysregulation, evasion of immune surveillance and therapy resistance (Hanahan and Weinberg, 2011; Masuda and Izpisua Belmonte, 2013). The CCI network can evolve in time and promote tumor progression. Factors secreted by the primary tumor can alter the microenvironment of distant organs, preparing them for metastatic invasion. Endothelial cells supply tumors with nutrition and oxygen and enable cancer cells to metastasize to distant sites through blood and lymphatic vessels. The intercellular communication between these TME components and cells is a driver of cancer progression and significantly impacts the efficacy of therapeutic interventions (Su et al., 2024). A well-studied example of TME interactions involves inflammatory cancer-associated fibroblasts (CAFs), which interact with endothelial cells to promote angiogenesis and tumor proliferation in bladder cancer and are a primary source of pro-angiogenic factors such as VEGF or PDGF (Su et al., 2024). Another well-studied example of TME interaction is the one with Tumor Associated Macrophages (TAMs) (Balaban and Cohen, 2025). Tumor cells produce chemokines to recruit circulating monocytes, which then differentiate into macrophages via tumor-secreted factors and ultimately establish a specific TAM phenotype via polarization signals. TAMs in turn favor the tumor by secreting growth factors, remodeling the ECM, inducing epithelial-mesenchymal transition (E-M transition) and suppressing immune responses (Balaban and Cohen, 2025). An emerging component of the TME is constituted by innervating neuronal cells, which might promote tumor growth through a feedforward loop involving the crosstalk between NGF and GPCR signaling in pancreatic and gastric cancers (Renz et al., 2018; Hayakawa et al., 2017).

The study of the TME involves the analysis of cell communication networks to identify interacting cell types, their molecular mechanisms and therapeutic targeting opportunities (Baghban et al., 2020). Early draft networks of receptor ligand pairs mediating cell-cell communications among human primary cell types revealed an extensive signaling network mediated by multiple receptor-ligand axes across cell types, characterized by high cell-type specificity of plasma membrane and secreted proteins, and a younger evolutionary age than intracellular proteins, also showing that most receptors had evolved before their ligands (Ramilowski et al., 2015). The analysis of cell-cell signaling from gene expression measurements of either bulk or single-cell transcriptomics data sets has become a routine functional analysis through the availability of comprehensive proteinprotein interaction (PPI) databases and recent advances in RNA sequencing (RNA-seq) technologies (Armingol et al., 2021). With the advent of single cell and spatial transcriptomics techniques, several computational approaches have been developed to infer the network of CCIs based on gene expression information (Armingol et al., 2021, 2024). Mathematical models exploiting network motif analysis have also been proposed to describe the TME in terms of small circuits of interacting cell types (Mayer et al., 2023). The analysis highlighted a hierarchical network of cellular interactions, with CAFs at the top of the hierarchy secreting factors mainly to TAMs. Here we applied a network motif analysis approach to study CCI circuits in cancer bulk RNA-seq data, which we chose as they allow their profiling across large patient cohorts and different cancer types, establishing more robust correlations with patients’ clinical data. In particular, we analyzed CCIs from gene co-expression networks, previously shown to be able to effectively identify prognostic genes across tumors (Yang et al., 2014) as well as to be predictive of direct PPIs and stable complexes (Miglionico et al., 2024), and demonstrated how CCI circuits can be exploited both as prognostic biomarkers as well as for immune response prediction.

## 3 Results

### 3.1 Cell-Cell interactions are highly co-expressed in cancer and highly co-evolved

We analyzed the expression of receptor-ligand pairs from large scale bulk transcriptomics data from 47 healthy and 32 cancer tissues, for a total of 15 matched tissues from over 15k unique samples (see Methods). We detected a similar number of genes despite tumor tissues generally having more samples than normal ones (Supplementary Figure 1A,B). We assessed gene co-expression individually in each healthy and cancer tissue using the weighted gene correlation network analysis (WGCNA) method (Langfelder and Horvath, 2008) and identified tissue-specific clusters. Both the number and sizes of clusters in each tissue were comparable across conditions (Supplementary Figure 1C,D). We investigated receptor-ligand pairs involved in mediating cell-cell communication in the gene co-expression network. We used a reference database of receptor-ligand interactions obtained by integrating multiple resources (see Methods), for a total of approximately 4k interactions. We found 2794 CCIs co-expressed within the same cluster in at least one healthy tissue, and 2301 in at least one cancer tissue (Supplementary Table 1). Despite detecting more unique co-expressed interactions in normal tissues (Supplementary Figure 1E), a comparable number of CCIs have their genes in the same co-expression network’s modules in tumor and normal tissues (Supplementary Figure 1F), indicating that CCIs are significantly more likely to cluster together in tumor tissues (*p* = 1.3 *·* 10*^−^*^87^; Figure 1A). We found that co-expression in cancer tissues is overall more predictive of receptor-ligand interactions than co-expression in healthy tissues, as the distribution of AUROC values obtained for this classification task using co-expression was significantly higher for cancer tissues than normal tissues (*p* = 7.7 *·* 10*^−^*^11^; Figure 1B). We found many CCIs that are exclusively co-expressed in the same cluster in healthy tissues and never in cancer, e.g. ACE2-AGTR1 or VEGFD-FLT4 (Figure 1C), likely representing specific, physiologically-relevant interactions that are dysregulated in tumors. Other interactions, such as EFNA4-EPHA1 or LTA4H-LTB4R are found more recurrently co-expressed in healthy tissues and, to a lower extent, in cancer ones (Figure 1C). We also found widespread CCIs recurrently co-expressed in the same cluster across cancer tissues (Figure 1C). The majority of CCIs recurrently co-expressed in cancer (e.g. SELPLG-SELL, HLA-DOA1-CD4, CCL5-CCR5 or CSF1-CSF1R) are also observed in healthy tissues, although less frequently, suggesting that these interactions might be co-opted in an aberrant way by the tumor (Figure 1C). A total of 236 interactions are exclusively co-expressed in tumor tissues, including the CXCL10-CXCR3 and CXCL11-CXCR3 axes (Figure 1C; Supplementary Table 1), which have recently emerged as major players in tumor growth and metastasis (Tokunaga et al., 2018).

**Figure 1:**
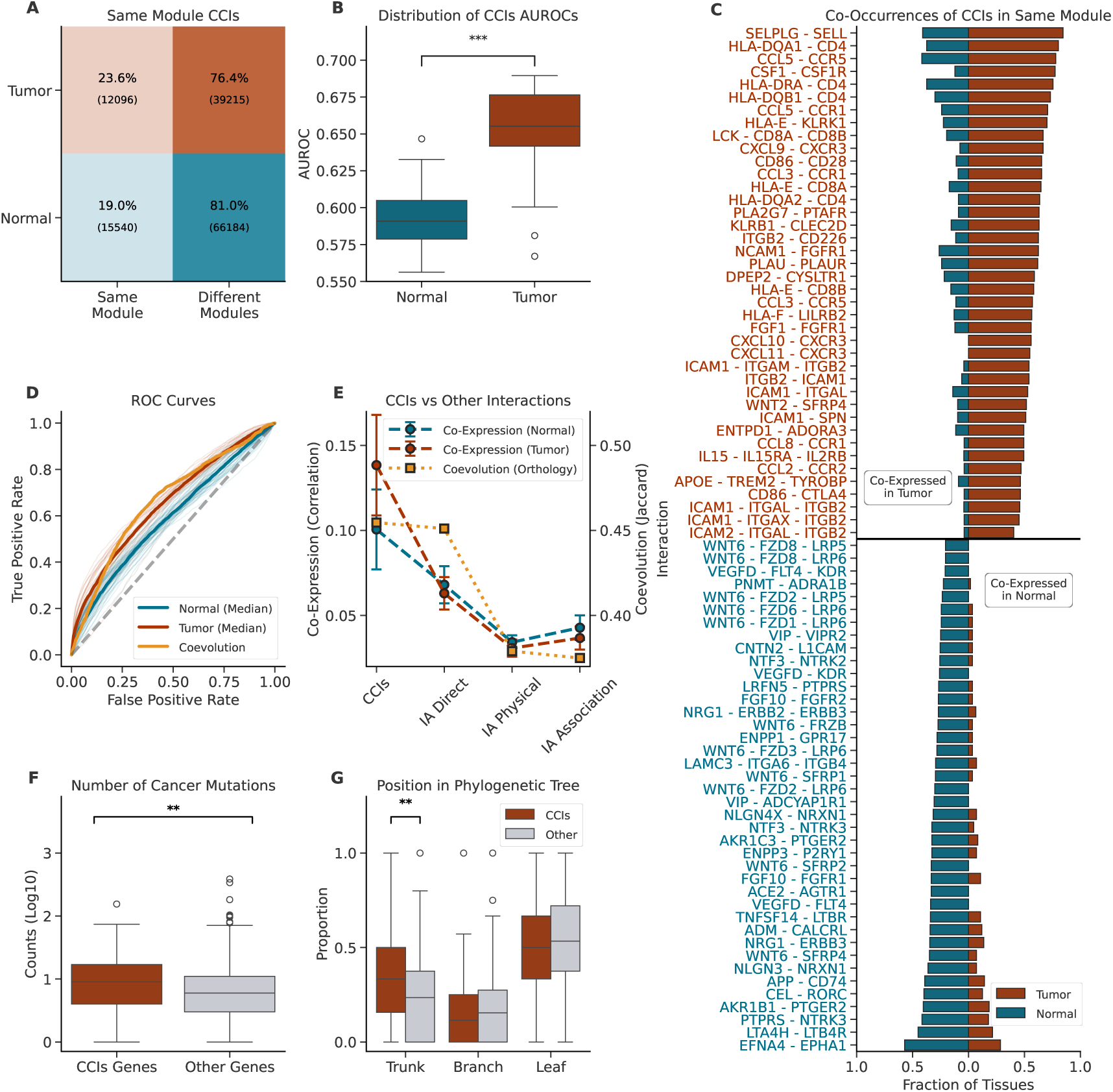
Co-expression and coevolution of CCIs. (A) Contingency table representing the co-occurrence of CCI genes in the same module in tumor and normal tissues; (B) Distribution of AUROCs (y-axis) in tumor and normal tissues (x-axis) for the classification task of discriminating gene pairs involved in CCIs from all other gene pairs based on co-expression; (C) Bar plot of the frequency of co-occurrence of CCI genes in the same module in tumor (red) and normal (blue) tissues, only the top 20 most commonly co-expressed interactions in tumor or normal tissues are shown; (D) ROC curves of individual (semi-transparent lines) tumor (red) and normal (blue) tissues and medians (opaque lines) across all tumor and normal tissues. The orange curve is obtained using orthology-based coevolution; (E) Average values of co-expression (left y-axis) and co-evolution (right y-axis) for CCIs and other interaction types from IntAct (x-axis). For co-expression, markers and error bars represent the mean and standard deviation of the in-tissue averages for tumor (red) and normal (blue) conditions; (F) Distribution of total number of mutations observed in CancerTracer (y-axis) for CCI genes (red) and other genes (gray). Only genes appearing in COSMIC’s Cancer Gene Census are considered; (G) Distributions of the proportion of the total number of mutations observed in different parts of the phylogenetic tree (x-axis) for CCI genes (red) and other genes (gray). Only genes in the Cancer Gene Census are considered. Significance is assessed using Mann-Whitney U test (* *p <* 0.05, ** *p <* 0.01, *** *p <* 0.001, BH corrected, two-tailed).

Co-evolution, assessed as the co-presence of gene pairs in sequenced genomes (see Methods), is also highly informative about CCIs and outperforms in predictiveness tumor co-expression (Figure 1D), suggesting that the strengthening of the co-expression of CCIs might mirror an evolutionary process. Moreover, we compared the co-expression patterns of CCIs with those of other known PPI types from a human reference interactome (i.e. IntAct (Del Toro et al., 2022)). Notably, co-expression of receptor ligand pairs is significantly higher compared to other IntAct interaction types (i.e. *Direct* interactions, *Physical* interactions or *Associations*), considering either healthy or cancer tissues (Figure 1E). Particularly in tumor, CCIs have a highly significant greater co-expression with respect to direct interactions (tumor *p* = 8.6 *·* 10*^−^*^12^, normal *p* = 2.2 *·* 10*^−^*^10^; Figure 1E). Additionally, CCIs are the interaction class showing the greatest increase in co-expression from healthy to cancer tissues (Figure 1E). CCIs, and to a lesser extent *IntAct Direct* interactions, are also characterized by higher co-evolution compared to interaction pairs from other IntAct classes (Figure 1E). To learn more about the evolutionary dynamic patterns involving genes mediating cell-cell communication, we also investigated the somatic mutations burden of cancer driver genes (from the Cancer Gene Census (Sondka et al., 2018)), distinguishing between genes involved in cell-cell communication (93 unique genes) and others (623 unique genes). We considered a curated database of cancer clonal evolution and heterogeneity (Wang et al., 2020), to stratify somatic mutations based on whether they affect the trunk, the branches or the leaves of the cancer phylogenetic tree, determined according to their emergence during tumor evolution (see Methods). Interestingly, we found that driver genes involved in cell-cell communication are significantly more mutated than other cancer driver genes, both overall as well as considering trunk mutations only (Figure 1F,G). On the other hand, we observed a tendency for higher burden of later-stage mutations (branch and leaf mutations) on other cancer driver genes (Figure 1G). Analysis of somatic mutations on cancer driver genes suggests that CCI genes are mutated earlier, suggesting an evolutionary pressure that determines their early fixation during tumor evolution.

### 3.2 Tumor mirrors natural evolution of GPCR signaling axes

We further investigated the different extent of co-expression and co-evolution pressures exerted by tumorigenic processes on receptor-ligand interactions, compared to intracellular interactions. We considered G protein Coupled Receptors (GPCRs), the largest family of transmembrane receptors, as a prototypical example of a receptor family whose members mediate interactions both with extracellular ligands and with intracellular transducers, e.g. heterotrimeric G proteins. Hence, this is an ideal system to monitor the different extents of regulatory pressure and evolutionary dynamics exerted from the extracellular and intracellular environments converging on the transmembrane receptor proteins. We compared the degree of co-expression of extracellular ligand-GPCR pairs and intracellular GPCR-G Protein pairs in healthy and cancer tissues. We found out that, while in healthy tissues ligands and G proteins have almost comparable levels of co-expression with cognate receptors (Kolmogorov-Smirnov distance *D* = 0.02, *p* = 3.8 *·* 10*^−^*^03^, Figure 2A), in cancer the ligand-GPCR pairs are significantly more co-expressed than GPCR-G Proteins pairs (*D* = 0.06, *p* = 4.1 *·* 10*^−^*^18^; Figure 2B). Likewise, natural co-evolution with the cognate receptor is significantly higher for ligands than G proteins (*D* = 0.20, *p* = 1.1 *·* 10*^−^*^15^; Figure 2C). We also correlated the natural co-evolution with the co-expression by averaging the values of GPCRs with their respective ligands, as well as the values relative to ligands and their respective GPCRs, for each tissue. We observed that highly co-expressed ligands and GPCRs are also characterized by stronger co-evolution scores in the vast majority of cancer tissues (Supplementary Table 2). Specifically, significant positive correlations were observed for ligand co-expression and coevolution in 24 tumor types (Pearson correlation, *p <* 0.05), including kidney (Supplementary Figure 2A) and breast tumors (Supplementary Figure 2B). Among these, five tumor types—such as breast cancer (Supplementary Figure 2C)—also showed significant associations for GPCRs. In contrast, no significant correlations were detected between co-expression and coevolution for either ligands or GPCRs in any of the healthy tissues analyzed (Supplementary Figures 2D,E,F). We also evaluated the overlaps of patients with both ligands and GPCRs mutated or both G proteins and GPCRs mutated in different parts of the phylogenetic tree. Despite CCIs’ tendency to mutate individual genes early (Figure 1G), we found no significant difference between the GPCR’s intracellular and extracellular sides in the earlier stages of tumor progression (trunk and branches, Figures 2D, E) when considering pairs of interacting genes. However, co-occurrences of mutations affecting ligands and their corresponding GPCR become significantly higher than mutations affecting G proteins and their receptors, when the whole phylogenetic tree is considered (*D* = 0.11, *p* = 5.1 *·* 10*^−^*^5^, Figure 2F), indicating that mutations affecting GPCRs are more frequently associated with mutations affecting the cognate ligand, rather than cognate G Protein. This effect, however, is not observed in the earlier stages of tumor progression, suggesting that a stronger selective pressure on the extracellular side of the cell membrane might emerge as the tumor evolves. Taken together, these analyses confirm also for GPCRs a higher co-expression of CCIs mediated on the extracellular side of the tumors and its TME, which is mirrored by the stronger evolutionary pressure observed in both natural and cancer-related evolution. Moreover, phylogenetic analysis of cancer evolution reveals that the stronger tumor coevolution on the extracellular side is characteristic of later stages of tumor evolution.

**Figure 2:**
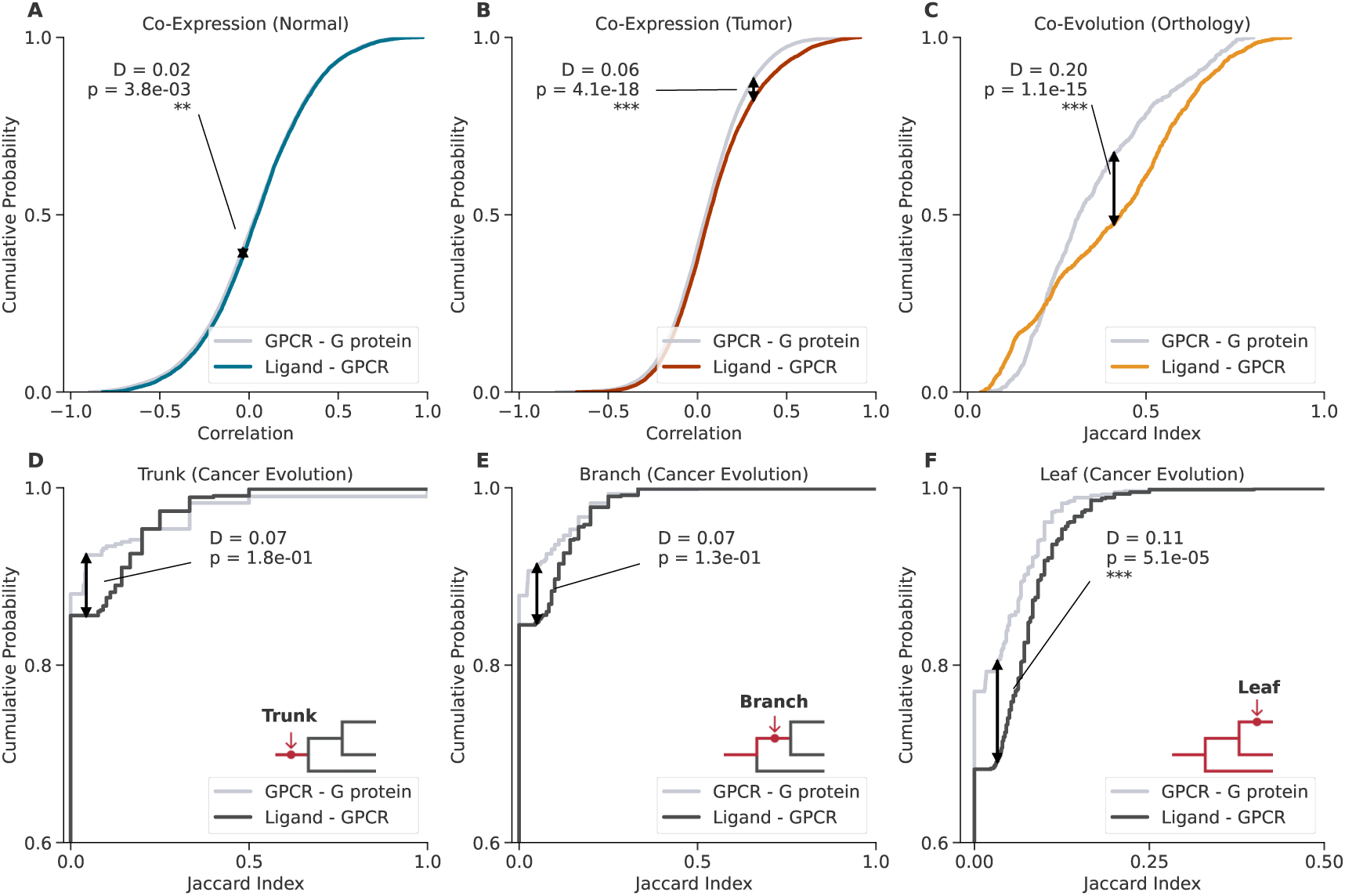
Cumulative distributions of co-expression and coevolution for GPCR-G protein and ligand-GPCR pairs. (A - B) Co-Expression values; (C) Orthology-Based coevolution; (D) Cancer coevolution of trunk mutations; (E) Cancer coevolution of mutations up to branch (trunk and branch mutations only); (F) Cancer coevolution of the whole phylogenetic tree (trunk, branch and leaf). Vertical black arrows represent the maximum distance (D statistics) and p-values refer to 2-sample Kolmogorov-Smirnov tests comparing the GPCR-G protein and ligand-GPCR distributions (* *p <* 0.05, ** *p <* 0.01, *** *p <* 0.001, BH corrected, two-tailed). In cancer coevolution plots, diagrams are added to illustrate the portion of the phylogenetic tree under consideration.

### 3.3 Major rewiring of ligand-receptor interactions in cancer

We analyzed the centrality properties of the CCI component of the co-expression network (see Methods). Healthy tissues display higher diversity, with more unique hub genes, which are tissue specific and rarely conserved across tissues (Figure 3A). On the other hand, in cancer there are overall fewer hub genes, which display higher conservation across tissues (Figure 3A). Indeed, while hub genes in healthy tissues are conserved at most in 40% of tissues (Figure 3A,B), in cancer they are shared in more than 90% of tissues (Figure 3A,C). This trend is possibly reinforced for genes involved in cell-cell communication, whose prevalence, relative to other hub genes, gets higher when considering genes shared by multiple cancer tissues. For instance, around one third of hub genes shared by at least 80% of cancer tissues are involved in CCIs (Figure 3C). This result suggests that healthy tissues have more specific transcriptional programs, while cancer tissues lose such specificity, being characterized by more similar transcriptional programs. Genes mediating cell-cell communication are highly conserved in the co-expression networks, suggesting their relevance for the interaction of the cancer cell with the TME. We also considered the biological function of hub genes by computing the connection strength contributed by hub genes in each biological process (MSigDB hallmark gene set collection (Liberzon et al., 2015); see Methods). This revealed that the processes that are more tightly connected to hub genes in tumor tissues are *Allograft Rejection*, *Inflammatory Response* and *KRAS Signaling Up*, all of which display significantly higher connectivity in cancer tissues than in normal ones (Figure 3D). Other processes, like *Adipogenesis*, *Glycolysis* and *P53 Pathway*, exhibit reduced connectivity with cancer hub genes relative to their connectivity in normal tissues (Figure 3D). Overall, 20 processes have significantly (*p <* 0.05) higher connectivity in normal tissues, while only 11 have stronger connectivity in tumor tissues. We investigated the rewiring of ligand-receptor pairs in co-expression networks from healthy to cancer tissues, by checking whether genes of a given interacting pair display coordination with the expression profiles of the same modules. We evaluated the similarity of module membership of interacting gene pairs via cosine similarity (see Methods). We found out that CCI is the interaction type undergoing the greatest extent of rewiring in cancer with respect to healthy tissues, expressed as a significantly different distribution of average cosine similarities that are higher in cancer compared to healthy tissues (*p* = 0.01, Figure 4E), indicating a stronger tendency of interacting ligand-receptor pairs to coordinate with the same module in cancer. Other interaction types, such as *Direct*, *Physical* and *Association*, as defined in IntAct, all have a comparable distribution of cosine similarities between cancer and healthy tissues (Figure 3E). We also analyzed co-occurrences of interacting genes in the same module, finding that certain tissues display a prevalence of clustered CCIs in healthy tissues (e.g. liver, skin, uterus), while others in cancer (e.g. prostate, esophagus and kidney) (Figure 3F). For example, in the prostate a large pool of CCIs that are uncorrelated in the healthy tissue clustered together in prostate tumors, mainly contributing to the largest cluster and, to a lesser extent, to smaller clusters (Figure 3G). These CCIs are enriched for processes such as *E-M Transition*, *Inflammatory Response* and *Angiogenesis* (Supplementary Figure 3A). Very few CCIs that are clustered in the healthy prostate become uncorrelated in cancer (Figure 3G) but, interestingly, this set of cancer-disrupted CCIs is also enriched for *E-M Transition* (Supplementary Figure 3A). In contrast, the liver – despite sharing some enriched processes with the prostate (Supplementary Figure 3B) – predominantly displays clustered CCIs in the healthy tissue (Supplementary Figure 3C), with the biggest tumor clusters formed by a combination of both previously clustered and unclustered CCIs (Supplementary Figure 3C). To illustrate how the cancer rewiring process affects biological processes, we generated network visualizations of genes involved in *E-M Transition* in the prostate (Figure 3H) and the liver (Supplementary Figure 3D). These visualizations highlight how both the activation of new CCI axes and the disruption of existing ones contribute to reshaping the co-expression networks. In particular, in the prostate the increased tumor co-expression leads to the emergence of three distinct cancer modules (Figure 3H), while in the liver, two modules of the normal tissue are disrupted in cancer (Supplementary Figure 3D).

**Figure 3:**
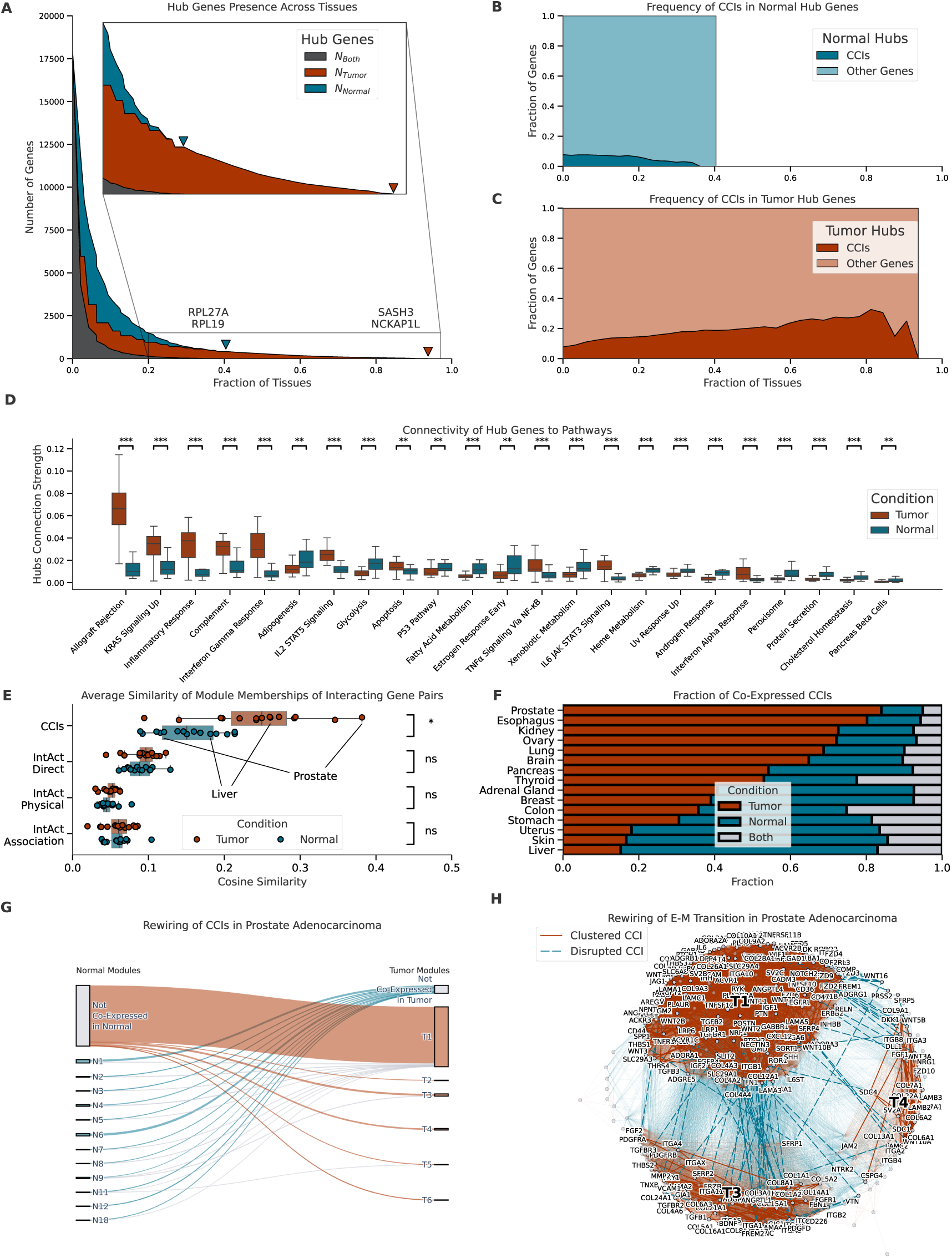
Rewiring of hubs and modules in co-expression networks. (A) Presence of hub genes across multiple tissues. For each fraction of tissues (x-axis), the plot shows the number of genes (y-axis) that are hubs in at least that fraction of tumor tissues (red), normal tissues (blue) or both normal and tumor tissues (gray). Triangles indicate the maximum fraction observed in normal and tumor hubs, and labels indicate the corresponding genes. The inset box shows a zoomed view of the highlighted region; (B) Fraction of hub genes involved in CCIs (y-axis) vs fraction of normal tissues the genes are hubs in (x-axis); (C) Fraction of hub genes involved in CCIs (y-axis) vs fraction of tumor tissues the genes are hubs in (x-axis); (D) Distributions of connection strengths (y-axis) of hub genes to genes involved in biological processes from MSigDB Hallmarks gene sets (x-axis) in tumor (red) and normal (blue) tissues. Significance bars represent Mann-Whitney U tests (** *p <* 0.01, *** *p <* 0.001, BH corrected, two-tailed, *p >* 0.01 not shown); (E) Distribution of average cosine similarity of module membership vectors (x-axis) for interacting gene pairs (CCIs and other interaction types from IntAct, y-axis) across tumor and normal tissues. Only tissues for which both tumor and normal samples are present are considered. Significance bars represent the results of Wilcoxon signed-rank tests (ns *p >* 0.05, * *p <* 0.05, BH corrected, two-tailed); (F) Fraction of CCIs with all genes located in the same module under tumor, normal, or both conditions. CCIs that are not co-expressed in either tumor or normal are excluded; (G) Sankey diagrams describing changes in module memberships between healthy prostate and prostate adenocarcinoma. Normal modules (blue, left side) and tumor modules (red, right side) are ranked based on their size. The height of each module is proportional to the number of co-expressed CCIs in it. CCIs that are co-expressed in only one condition are highlighted in blue (same module in normal, different modules in tumor) or red (same module in tumor, different modules in normal). CCIs that are not co-expressed in any condition are not shown. The remaining CCIs (gray) are co-expressed in both conditions; (H) Network visualization of *E-M Transition* in prostate. Only genes involved in the process and CCIs that connect to them are shown. CCIs that are co-expressed in the same module only in cancer (only in normal) are highlighted in red (blue). Links that don’t represent CCIs are semi-transparent. Labels highlighting tumor modules T1, T3 and T4 are positioned corresponding to the centroid of the respective clusters.

**Figure 4:**
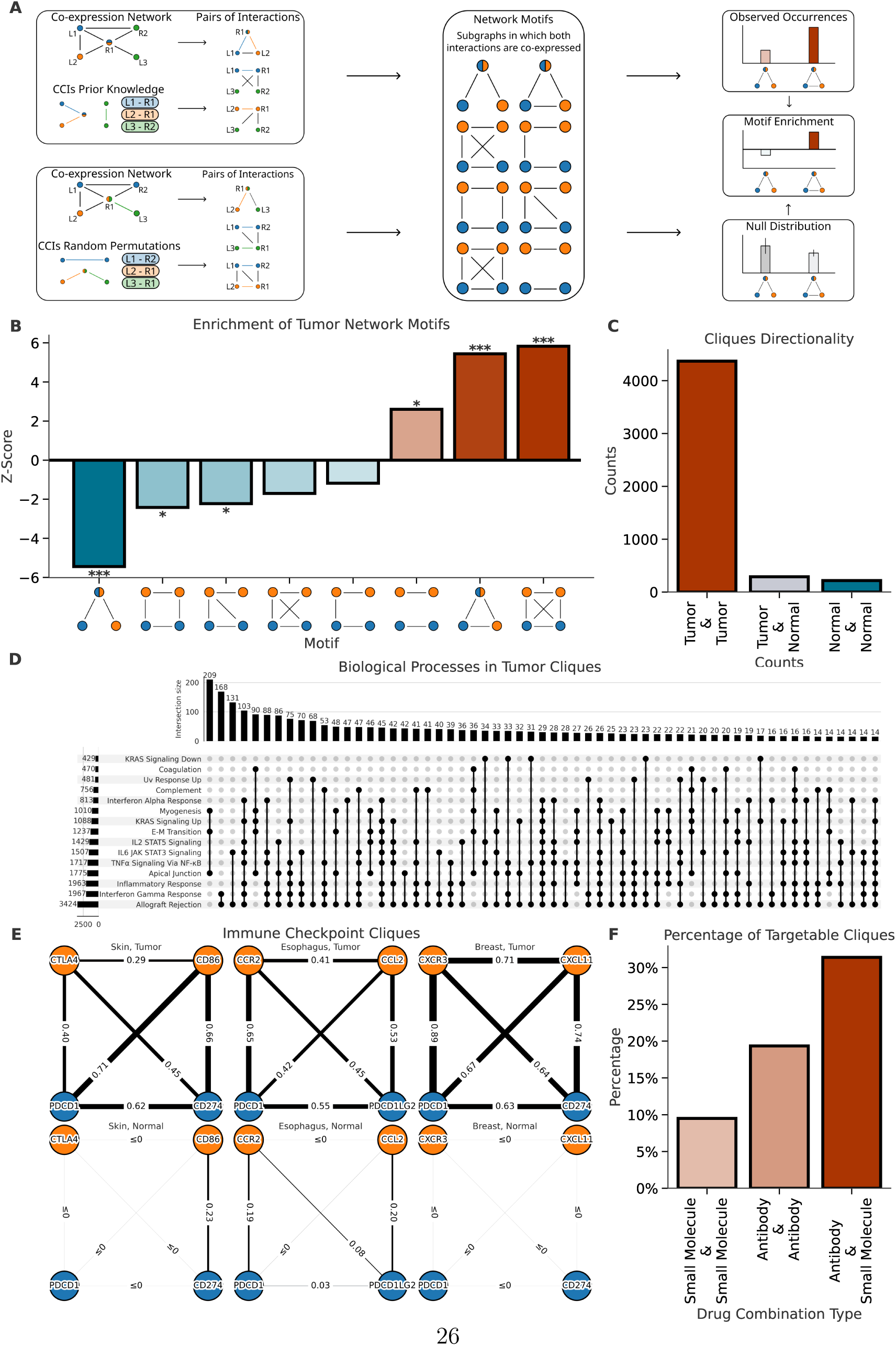
CCI circuits. (A) Overview of network motifs analysis. Individual tissues’ co-expression networks are merged into a single network (see Methods). All pairs of co-expressed CCIs are sorted into possible network motifs and the procedure is repeated for random permutations of the co-expression network. The observed occurrences of each motif are compared to the background distribution of permutations, obtaining a Z-score representing its enrichment; (B) Z-scores (y-axis) represent the enrichment of each network motif (x-axis) in tumor co-expression networks, where edges indicate higher co-expression in tumor versus normal samples. p-values are derived from permutation testing (* *p <* 0.05, ** *p <* 0.01, *** *p <* 0.001, BH corrected, two-tailed); (C) Bar plot of the number of cliques (links represent any difference between conditions) where both CCIs show higher co-expression in tumor, normal, or with one CCI more co-expressed in tumor and the other more co-expressed in normal; (D) UpSet plot describing the membership of genes of tumor cliques (higher co-expression in tumor) to biological processes (MSigDB Hallmarks); (E) Co-expression values of (left) genes involved in the CTLA4-CD86 & PDCD1-CD274 clique in tumor and normal skin, (center) CCR2-CCL2 & PDCD1-PDCD1L2 clique in esophagus, CXCR3-CXCL11 & PDCD1-CD274 in breast. (F) Bar plot of the percentage of clique circuits have both CCIs targeted by at least one approved combination of antibodies, small molecules or both.

### 3.4 CCIs form co-expressed circuits in cancer suggesting crosstalk among different cell-cell communication pathways

We investigated the emergence of circuits of two distinct CCIs (two-CCIs circuits) displaying coordinated expression, possibly indicating crosstalk of two different signaling axes. These crosstalk mechanisms could belong to either the same or different pathways and might happen in either an autocrine or paracrine fashion. We aggregated individual tumor and normal co-expression networks into a single network where links connect pairs of genes that display differences in co-expression between the two conditions (see Methods). We enumerate all possible pairs of connected CCIs, and selected all subgraphs made of two CCIs, which might involve either three nodes (e.g. two ligands binding to the same receptor) or four nodes (two ligand-receptor pairs). Each two-CCIs circuit can form one of 8 possible network motifs (2 triads and 6 tetrads) based on the configuration of the links connecting its genes (Figure 4A). We compared the frequency of each network motif with a background distribution obtained by randomly shuffling ligand-receptor interactions (see Methods). We used z-scores to represent enrichment (z > 0) or depletion (z < 0) of each network motif. First, we considered links indicating greater co-expression in tumors. In the tumor case, the only significantly enriched motifs are either characterized by disjoint CCIs (*p <* 0.05), or, conversely, those in which each possible pair of genes is co-expressed, forming a fully connected triad or tetrad clique (*p <* 0.001, Figure 4B, Supplementary Table 3). This suggests that cancer favors crosstalk between CCIs whose components are all co-expressed with one another, i.e. any possible pair of genes within the clique is co-expressed. Any possible intermediate form of crosstalk, where only a fraction of participating genes are co-expressed, happens less frequently than expected (Figure 4B). We strengthened this observation by repeating the procedure while also considering links that indicate lower co-expression in tumors and classifying them based on the condition in which the 2 CCIs involved were more strongly co-expressed. We found that the vast majority of the cliques connect two CCIs whose co-expression increases in cancer, followed by cliques in which increased co-expression of a CCI corresponds to decreased co-expression of the other one and, finally, by cliques whose CCIs are concordantly more co-expressed in healthy tissues (Figure 4C). We hypothesize that cliques in which all links concordantly indicate increased cancer co-expression might underline convergent mechanisms between distinct signaling axes in tumor settings and focus on only those circuits for the rest of the analyses. We inspected the processes that are more frequently co-represented within two-CCIs clique circuits. We found that the combination of CCIs involved in *Allograft rejection* with *Interferon Gamma Response* is the most frequent one, being represented in 166 cliques (Figure 4D, Supplementary Figure 4). Other highly recurrently crosstalking processes are *Apical junction* with *E-M Transition* and *Myogenesis*, co-present in 119 tetrad cliques, and Allograft rejection with *IL6 JAK STAT3 signaling*, co-present in 109 tetrad cliques (Figure 4D). Overall, *Allograft rejection* is the single process most frequently involved in crosstalk with other processes, followed by *Interferon Gamma Response* and *Inflammatory Response* (Figure 4D). Frequently observed genes within clique circuits include integrins like ITGB1, ITGA11 and ITGB2, T cell co-receptors genes CD4, CD8A and CD8B, chemokine receptors like CXCR3 and CCR5, and chemokines including CCL8, CCL5, and CXCL13, each participating in more than 250 distinct clique circuits (Supplementary Table 3). Interestingly, numerous cliques incorporate immune checkpoints such as CTLA4, which, along with its corresponding ligands, is observed in 239 distinct clique circuits, or PDCD1—involved in 128 cliques—often reflecting known examples of crosstalk between distinct CCIs in specific tissues. For instance, the CTLA4 and PDCD1 axes—remarkably coordinated in skin cancers (Figure 4E)—show increased response rate when co-targeted in metastatic melanoma Rotte (2019). The CCL2-CCR2 & PDCD1-PDCD1LG2 circuit, vastly co-expressed in esophageal cancers (Figure 4E), promotes immune evasion via TAM recruitment in esophageal squamous cell carcinoma mouse models (Qin et al., 2023). In breast cancer, PDCD1-CD274 coordinates with CXCL11-CXCR3 (Figure 4E), the latter influencing response to checkpoint inhibitor therapies targeting PDCD1 (Gomes-Santos et al., 2021). We also explored therapeutic targeting opportunities on two-CCIs circuits co-expressed in cancer. We have mapped known small molecules and antibodies targeting the receptors involved in two-CCIs clique circuits. We found that 36% of two-CCIs clique circuits have both signaling axes targeted by at least one approved drug (either small-molecule or antibody). The majority of two-CCIs cliques are targeted by combinations of small molecules drugs and antibodies, followed by cases where both receptor axes are targeted by antibodies and those targeted by small-molecules only (Figure 4F). However, most tetrad cliques (64%) are not yet targeted by combinations of approved therapies, suggesting an enormous space for discovery of new therapeutic approaches.

### 3.5 CCIs circuits are associated with patient survival

We considered whether two-CCIs cliques circuits could represent functional units that synergize to promote specific patient phenotypes, such as survival. In this respect, we investigated the use of expression of two-CCIs circuits to stratify patients based on survival times. We identified two-CCIs circuits whose expression in patient samples showed a statistically significant association with patient survival. We then filtered only those circuits whose expression led to better survival statistics and patient stratification compared to the expression of individual CCIs (see Methods). Intriguingly, we found an almost linear relationship (*r* = 0.87*, p* = 0.005) between network motif enrichment (z-score), and the likelihood of association with patient survival (Figure 5A). Remarkably, we found out that cliques (both tetrads and triads), that represent the most enriched network motifs, are also the most strongly associated with patient survival (Figure 5A). Overall, we identified 2283 two-CCIs clique circuits significantly associated with patient survival in at least one of the 29 cancer tissues considered for the survival analysis, with the highest number of such prognostic circuits observed in brain and skin cancers (Supplementary Table 4).

**Figure 5:**
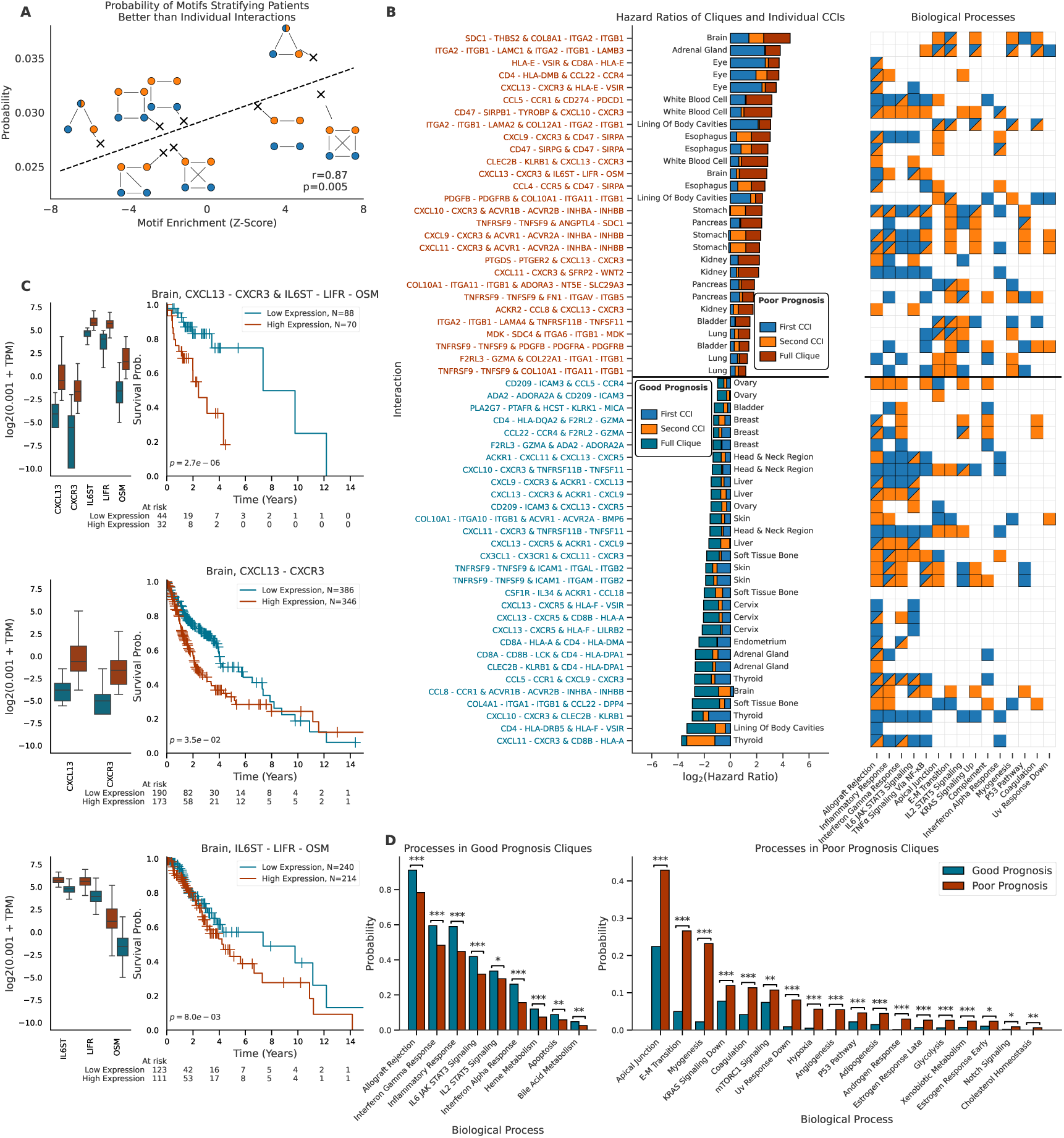
Crosstalking interactions are associated with patient survival. (A) Probability of each possible network motif of 2 CCIs improving patient stratification (y-axis) vs enrichment of the motif (x-axis); (B) Barplot of cliques hazard-ratios (left) and heatmap of associated biological processes (right). Each clique has three stacked bars: the blue bar shows the log hazard ratio of the first CCI. The orange bar shows the increase when using the second CCI (labels are arranged so that the second CCI always offers better stratification). The third bar (colored like the label) shows the added value of using the full clique over individual CCIs. Label colors indicate poor prognosis (red, positive log HR), or good prognosis (blue, negative log HR). The heatmap shows which biological processes include genes from the first CCI (blue), second CCI (orange), or both. Top 3 cliques per tissue by rank aggregate (Borda count) of c-index and increase of hazard-ratio with respect to individual CCIs are shown; (C) Expression values and Kaplan-Meier curves with patient stratification based on expression of the full CXCL13-CXCR3 & IL6ST-LIFR-OSM clique (top), expression of the CXCL13-CXCR3 CCI only (center) or expression of the IL6ST-LIFR-OSM CCI only (bottom) in brain cancers. p-values refer to logrank tests; (D) Probability (y-axis) of good prognosis (blue) and bad prognosis (red) cliques containing genes from each biological process (x-axis). Only processes that are significantly more likely to appear in good prognosis cliques (left) or poor prognosis cliques (right) are shown (Fisher’s exact test, * *p <* 0.05, ** *p <* 0.01, *** *p <* 0.001, BH corrected, two-tailed).

Several cliques involving interactions mediated by collagen and integrin are recurrently associated with poor prognosis. For example, in brain tumors, the clique formed by COL8A1-ITGA2-ITGB1 & SDC1-THBS2 is linked to shorter survival (Figure 5B). Similarly, in adrenal gland tumors, the ITGA2-ITGB1-LAMC1 & ITGA2-ITGB1-LAMB3 circuit is also associated with reduced survival (Figure 5B). These two-CCIs circuits are examples of cliques whose expression is mainly associated with lower survival and connect processes associated with *Apical Junction*, *E-M Transition*, *KRAS Signaling Up*, *Myogenesis* or *Coagulation*. Other clique circuits associated with poorer prognosis, such as CXCL13-CXCR3 & HLA–E-VSIR in uveal melanoma, CCL5-CCR1 & CD274-PDCD1 in leukaemia, CXCL13-CXCR3 & IL6ST-LIFR-OSM in brain tumors (Figure 5C) or CCL4-CCR5 & CD47-SIRPA in esophageal cancer mediate crosstalk between *Allograft rejection*, *Inflammatory Response*, *IL6 JAK STAT3 Signaling* or *Interferon Gamma Response* (Figure 5B). Among lower survival cliques, we also found some connecting *Allograft rejection* and *Inflammatory Response* with *Apical Junction* or *E-M Transition*, such as CCL5-CCR1 & CD274-PDCD1 in leukemia or CXCL9-CXCR3 & CD47-SIRPA in esophageal cancer (Figure 5B).

On the other hand, we have found different cliques associated with good prognosis in specific tumor types, such as CXCL11-CXCR3 & CD8B–HLA-A in thyroid carcinoma or COL4A1-ITGA1-ITGB1 & CCL22-DPP4 in sarcoma (Figure 5B). Interestingly, some cliques can also be associated with good or poor prognosis depending on the specific type of cancer. For example, lung cancer patients with high expression of the EGFR-HBEGF & ITGA6-ITGB1-LAMA2 circuit are characterized by shorter survival times than those with lower expression (Supplementary Figure 5A). On the other hand, expression of the same circuit is associated with longer survival times for esophageal carcinoma patients (Supplementary figure 5B).

In general, we found that 22% of the cliques mapping to *Allograft rejection* are significantly associated with either good or poor prognosis depending on the cancer tissue considered, while only 4% of the cliques mapping to *E-M Transition* are either favourable or unfavourable depending on the tissue considered (Supplementary Figure 5C). Overall, cliques associated with higher survival are enriched (*p <* 0.05) for processes such as *Allograft Rejection*, *Interferon Gamma Response*, *Inflammatory Response*, *IL6 JAK STAT3 Signaling* or *TNF Signaling via NF-kB* (Figure 5D). In contrast, cliques associated with lower survival display a drastic enrichment of processes such as *Apical Junction*, *E-M Transition*, *Myogenesis*, *Hypoxia* or *Angiogenesis* (Figure 5D). The near-exclusive involvement of these processes in cliques associated with lower patient survival indicates that their modulation leads unequivocally to poorer prognosis. On the other hand, processes such as *Allograft Rejection* or *Inflammatory Response* are more prevalent in cliques associated to higher survival rates, but they also appear, although to a lower extent, in poor-prognosis cliques, suggesting a more nuanced and context-dependent involvement of these interactions and processes in determining patient prognosis.

### 3.6 CCIs circuits are predictive of response to immunotherapy

We investigated the predictive power of CCI cicuits expression related to patient response to immunotherapy. We considered an integrated dataset of 1246 cancer patients, spanning 8 tissues, who underwent immunotherapy treatment (see Methods). We trained a machine learning predictor for immunotherapy response based on patient transcriptomics data (see Methods). We explored multiple expression signatures as distinct feature sets, including the entire transcriptome, all the genes participating in CCIs, the set of all genes appearing in two-CCIs clique circuits, and known immunotherapy response signatures collected from the literature (i.e. TIGER database (Chen et al., 2023)). For each of these feature sets, we trained both tissue-specific and pan-cancer tissue predictive models, which we evaluated on held-out testing sets in a stratified k-fold procedure (see Methods). We showed that models trained on genes involved in two-CCIs clique circuits performed better across tissues than those based on any other gene signature, including specialized immunotherapy response signatures such as *Tertiary Lymphoid Structures* (TLS) or *T Cell-inflamed Gene Expression Profile* (T Cell-inflamed GEP) (Figure 6A,B). Notably, they also outperformed models using the whole transcriptome in 5 out of 8 tissues when considering AUROC (*p* = 0.03; Figure 6A,B,C) (all tissues except skin, brain and kidney), and in 6 out of 8 tissues when considering AUPRC (*p* = 0.03, Figure 6C) (all except skin and brain), obtaining the best predictive performances in lung tumors (AUROC = 0.86, AUPRC = 0.79, Figure 6A, Supplementary Figures 6A,B). Similarly to survival analysis, we also trained models based on expression signatures of individual two-CCIs circuits and compared their performances with models trained on individual CCIs (Supplementary Table 6), finding that the tetrad clique is the network motif exhibiting the strongest predictive power (Supplementary Figure 6C). Crucially, several clique circuits hold high predictive power in specific cancer tissues and generate better predictions than those based on any of the individual CCIs that make the clique up (e.g. ACKR1-CXCL9 & CCL5-CCL4 in bladder cancer; Figure 6D; Supplementary Figure 6D). Notably, two-CCIs clique circuits performed significantly better than models trained on either random pairs of CCIs regardless of their co-expression, as well as models trained on other non-clique two-CCIs circuits (*p <* 0.001; Figure 6E). The predictive clique circuits are distinguished by a strong enrichment in *Allograft Rejection* (Figure 6F), underscoring the centrality of this process in mediating the response to immunotherapy, and by a marked depletion in *E-M Transition* (Figure 6F). The individual cliques showing the best predictive performances across tumor types are almost invariably involved in *Allograft Rejection* and frequently crosstalk with processes such as *Inflammatory Response*, *Interferon Gamma Response*, *TNF Signaling via NF-kB*, *IL2 STAT5 Signaling*, *KRAS Signaling Up* or *IL6 JAk STAT3 Signaling* (Supplementary Figure 6D). We further investigated the depletion of CCI genes involved in *E-M Transition* (Figure 6F) by evaluating the predictive power of individual genes. For each gene, we computed the AUROC independently in each tissue by treating the gene’s expression level as a continuous predictor of the outcome. Interestingly many genes have predictive power in specific tissues like ureter, lung, lymphatic tissue and stomach (Supplementary Table 7), the majority of which are associated with AUROC < 0.5 indicating that their expression leads to reduced response rates.

**Figure 6:**
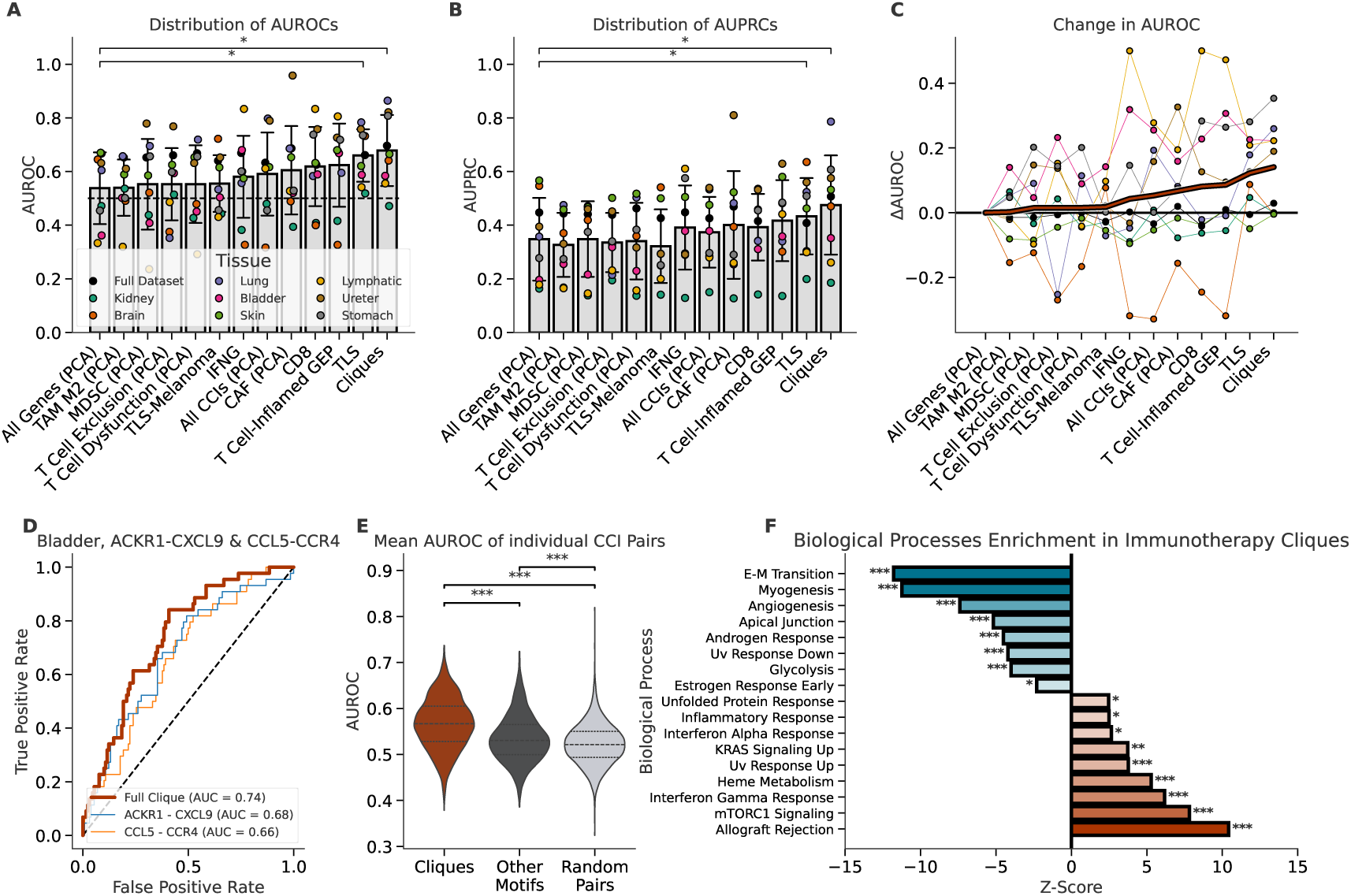
Crosstalking cliques are predictive of immunotherapy response. (A) Bar plot showing the AUROCs (y-axis) of different feature sets (x-axis); (B) Bar plots showing the AUPRCs (y-axis) of different feature sets (x-axis). Each point in the two bar plots represents a cohort, while bar heights and error bars indicate the mean and standard deviation. Feature sets are sorted by mean AUROC, and only those with mean AUROC higher than *All Genes (PCA)* are displayed. Significance was assessed using Wilcoxon signed-rank test, comparing each feature set against "All Genes (PCA)" (one-tailed); (C) Change in AUROC value for different feature sets. For each cohort, “All Genes (PCA)” is considered as a baseline with dots representing the increase/decrease in AUROC (y-axis) when changing features set (x-axis). The thick red line indicates the mean increase across all cohorts; (D) Example of ROC curves for models trained on the genes of the ACKR1-CXCL9 & CCL5-CCR4 clique (red), ACKR1-CXCL9 only (blue) and CCL5-CCR4 only (orange) for the bladder cohort.(E) Distribution of AUROCs (y-axis) for models trained on a single clique, any non-clique motif from the co-expression network or random pairs of CCIs (x-axis). Significance assessed with Mann-Whitney U test (BH corrected, two-tailed); (F) Enrichment (Z-score, x-axis) of biological processes (y-axis) in predictive cliques. Background distribution of occurrences of biological processes and p-values are obtained through permutation testing (BH corrected, two-tailed). (* *p <* 0.05, ** *p <* 0.01, *** *p <* 0.001).

## 4 Discussion

In this study we explored expression signatures and evolutionary dynamics of molecular interactions mediating cell-cell communication on large scale transcriptomics datasets from healthy tissues and cancer, and inspected their association with patient survival and response to immunotherapy. We found that CCIs are more co-expressed in cancer than in healthy tissues. Moreover, they are also more co-expressed when compared to other types of intracellular interactions within the same condition (i.e. both in cancer and in normal tissues). Here we show that CCIs are the class of interactions displaying the highest co-expression level, particularly in tumors, where receptor-ligand interactions exhibit greater coordination. Co-expression patterns are predictive of CCIs, in line with our previous work that shows that co-expression derived from cancer tissues is more predictive of proteins engaging in direct interactions or forming stable complexes (Miglionico et al., 2024). In contrast, normal tissues display a broader diversity of ligands and receptors, though their expression patterns are less tightly correlated. The stronger co-expression observed for CCIs compared to any other interaction class is also mirrored by a significantly higher co-evolution of the former when considering the co-presence of interacting partners in sequenced genomes. As a special receptor system, we considered GPCRs, since they evolved the fine-tuned capability to bind a set of intracellular transducers, i.e. heterotrimeric G proteins, upon activation mediated by the binding of specific extracellular ligands. This makes GPCRs the ideal receptor class to study the different extent of co-expression and co-evolutionary pressures acting on both sides of the cell membrane. The ligand binding pockets of GPCRs evolved to bind a huge variety of extracellular ligands, information which is crucial to guide the molecular phylogenetic classification of these receptor systems (Schiöth and Fredriksson, 2005). On the other hand, the binding interface of intracellular G protein is relatively more conserved, with sequence determinants of binding mainly residing on the G protein while the receptor displays less conserved and highly evolutionary dynamical determinants (Flock et al., 2017). Herein we show that at the gene-gene interaction level, GPCRs are more coevolved, in terms of co-presence within sequence genomes, with their ligands than with cognate G proteins. This trend correlates with higher co-expression of receptor-ligands pairs compared to receptor-transducer ones, especially in cancer. Overall, this suggests that the cancer transcriptional programs reinforce cell-cell communication, potentially providing an advantage for cancer cells to thrive within the TME. Such a process seems to mirror the natural evolution of CCIs which are characterized by a higher selective pressure compared to other categories of interactions. Along this line, we found significantly higher co-occurrences of somatic mutations at later stages of cancer evolution for receptor-ligand pairs compared to receptor-transducer pairs, supporting the notion of a higher selective pressure for CCIs also in cancer.

We have found a vast diversity of CCIs in healthy tissues, with certain interactions being exclusively associated with healthy tissues, potentially representing tumor-suppressive interactions, while other interactions are more prevalent in cancer tissues and might function as TME components. CCIs are central in co-expression networks, having greater hub character and being more conserved in cancer co-expression networks compared to other genes. On the other hand, healthy tissues display a greater diversity of co-expressed CCIs, which are less conserved, more specific and undergo a massive rewiring in cancer. The cancer rewiring process and the transcriptional program reinforcing CCIs also favors the coordination of specific circuits of multiple CCIs which tend to recurrently emerge in cancer networks. CCIs in these circuits are often fully-connected, forming cliques and possibly representing self-sustaining feedback-loops or indicating coordinated regulatory mechanisms establishing crosstalk between distinct processes exploited by cancer. Indeed, we found over-representation of certain processes such as *Allograft Rejection*, *Interferon Gamma Response*, *Inflammatory Response*, *Apical junction*, *E-M Transition*, *Myogenesis* or *IL6 JAK STAT3 Signaling*, which are frequently co-occurring in the same clique. Remarkably, we have found that the coordinated expression of many two-CCIs circuits is significantly associated with patient survival. Crucially, considering the expression of the whole two-CCIs circuits is often more significant than considering the expression of individual receptor-ligand interactions, especially when the circuit is a clique, suggesting a synergistic role of the clique participants in triggering processes contributing to a certain patient phenotype. In particular, the clique expression signatures associated with poor prognosis frequently participate in processes such as *E-M Transition* or *Myogenesis*, while others such as *Allograft Rejection* or *Inflammatory Response* might be associated with either lower or higher survival depending on the specific tissue and CCIs involved. In line with previous studies showing that CCI expression signatures (Lee et al., 2024; van Santvoort et al., 2025), or complex mutational signatures (Kong et al., 2024), could be harnessed to predict response to Immunotherapy, we showed that expression signatures based on two-CCIs clique circuits are more predictive of immunotherapy response, compared to the whole transcriptome, individual CCIs or other immunological expression signatures. Interestingly, the most enriched processes in predictive circuits are *Allograft rejection*, *mTORC1 Signaling* and *Interferon Gamma Signaling*, while the most depleted ones are *E-M Transition*, *Myogenesis* and *Angiogenesis*. This indicates that, although interactions involved in *E-M Transition*, *Myogenesis*, or *Angiogenesis* are clearly linked to poorer prognosis, their expression signatures are not able to specifically predict immunotherapy response. However, single *E-M Transition* genes contribute to the prediction of immunotherapy response in cancers with higher metastatic potential (lung, lymphatic tissue and stomach cancer) (Supplementary table 7): high cytokine expression in cancer cells (IL15) is associated to positive response to immunotherapy (as also observed in mouse (Hu et al., 2024)), whereas extracellular matrix, fibrosis and angiogenesis-related genes (collagen, PDGFRB, ITGB3) expressed in stromal cells are associated with resistance. Overall, the presence of multiple genes with different effects on immunotherapy response, the diversity in predictivity across cancer types and the small effect of most genes may explain why *E-M Transition* cliques have low predictive power. By focusing on more predictive processes, we can devise more universal markers. In particular, the expression of CCIs circuits involved in immune response processes such as *Allograft Rejection* and *Inflammatory Response* is associated with higher survival times and a higher predictive power for immunotherapy response. The expression signatures of two-CCIs clique circuits might represent new biomarkers to stratify patients and also open new avenues to synergistic targeted therapies. Many CCI axes are targeted by approved targeted drugs which could be used in combination with immunotherapies. For instance, we have found that a considerable amount of receptor-ligand pairs in two-CCIs cliques are targeted by a combination of small molecule drugs and antibodies, paving the way to the systematic exploitation of synergistic combination of targeted drugs with immunotherapies. The current analysis only considered bulk RNA-seq data. On one hand, the possibility to employ predictive models of either patient survival or immunotherapy based on CCI expression signatures derived from bulk RNA-seq data has undoubtedly the advantage to be more exploitable in clinical settings, given the higher cost-effectiveness of the methodology. Moreover it can also be combined with other approaches to predict targeted therapies based on bulk RNA-seq e.g. CellHit (Carli et al., 2025a,b). On the other hand, the current analysis lacks the information about TME cell type heterogeneity, which may limit its ability to capture more nuanced cell-cell interactions and specific cellular contributions to patient phenotype. Therefore, we plan to extend this analysis by exploring the role of CCI circuits associated with distinct TME components in large-scale, pan-cancer, single-cell RNA-seq atlases (Tyler et al., 2025).

## 5 Methods

### 5.1 Preprocessing

Bulk RNA-seq data from TCGA and GTEx was obtained from the UCSC Xena Browser (Goldman et al., 2020). Only primary tumor data was kept from TCGA. The resulting dataset includes 18305 samples, 7775 normal tissue samples from GTEx and 10530 primary tumor samples from TCGA. We first identified outliers based on quality control (QC) metrics. We only kept samples with the number of detected genes between 20K and 40K. Samples with less than 10 millions or more than 120 millions total counts were also removed, same for samples in which the top 100 most expressed genes accounted for more than 90% of total counts. The data were then split according to tissue and condition (e.g. normal liver and hepatocellular carcinoma were considered different tissues) and preprocessing was applied to each tissue. In particular, we considered the following QC metrics: logarithm of total counts, logarithm of genes with at least one count (adding a pseudocount to handle 0 values) and percentage of counts related to the top 100 most expressed genes. Samples in which any of the QC metrics was more than 5 median absolute deviations from the median value of the tissue were removed. To prevent the sparsity of the data from driving the formation of clusters of lowly expressed genes, we removed genes with less than 15 counts in more than 75% of the samples from each tissue. We normalized the data using DeSeq2 (Love et al., 2014) through PyDeSeq2 (v0.4.4) (Muzellec et al., 2023) and applied DeSeq2’s variance stabilizing transformation. To identify remaining outlier samples, we conducted principal component analysis (PCA) on the normalized data and retained the first 10 principal components. For each tissue group, the centroid in the PCA space was calculated, and samples located more than five standard deviations from their respective tissue centroid were removed. Tissue groups with fewer than 15 samples remaining after all QC and filtering steps were excluded from subsequent analyses.

### 5.2 Weighted gene correlation network analysis

In order to generate co-expression networks, we performed weighted gene correlation network analyses with the PyWGCNA (v2.0.0) (Langfelder and Horvath, 2008) python module (Rezaie et al., 2023). Co-expression networks were generated individually for each subtissue to avoid batch-effects (e.g., two different networks for sigmoid colon and transverse colon). We used the “signed hybrid” network type and set the threshold for the approximate scale-free criterion to *R*^2^ = 0.9. We named gene modules according to their ranking by size, as well as the condition of the tissue they belong to (N for normal, T for tumor). For instance, T1 is the name of the module containing most genes in a tumor tissue while the 5th module by size in a normal tissue is called N5. We used modularity (Newman, 2006) to assess that the modules actually reflect the community structure of the network. We found that 88% of the tissues display modularity *Q >* 0.3, and all have *Q >* 0.2 except for normal testis (Q = 0.05). Further inspection revealed lower number of modules and average connectivity than all other tissues, for this reason the normal testis co-expression network was deemed an outlier and excluded from the analysis.

### 5.3 List of interactions

The list of CCIs used in this work is obtained by merging CellPhoneDB (Efremova et al., 2020) and CellChatDB (Jin et al., 2021) (the latter is obtained from LIANA (Dimitrov et al., 2024)). Additionally, in the case of GPCRs, the list also includes biosynthetic enzymes as surrogates for GPCR organic ligands (Arora et al., 2024). For intracellular protein-protein interactions, we retrieved all interactions from the IntAct database (Del Toro et al., 2022) and grouped them into the following categories (in order of priority): *IntAct Direct* (both enzymatic and non-enzymatic), *IntAct Physical*, *IntAct Association*. Each interaction only appears in one category (e.g. an interaction that appears both in CCIs and *IntAct Direct* is removed from *IntAct Direct*).

### 5.4 Co-Expression

The following metrics were considered when investigating co-expression: Pearson correlation, adjacency (soft-thresholding) in WGCNA networks and co-presence in the same module. We classified each occurrence of a CCI in each tumor or normal tissue based on whether or not all the genes of the CCI are part of the same module and constructed a contingency table. Significance was assessed with fisher’s exact test. Gene pairs were then classified as either participating in cell-cell interactions (CCIs) or not, excluding pairs belonging to the same protein complex from the interacting category. ROC curves were generated using scikit-learn (v1.4.0) in order to assess the predictive power of adjacency in each tissue. Mann-Whitney U test (scipy v1.12.0) was used to compare tumor and normal AUROC values. We also considered PPIs from IntAct. For each type of interaction in our list (CCIs, *IntAct Direct*, *IntAct Physical*, *IntAct Association*) the mean co-expression (Pearson correlation) of interacting protein pairs in each tissue was evaluated. Mann-Whitney U tests were employed to compare the distributions associated with different interaction types (e.g. CCIs vs *IntAct Direct*). P-Values (two-tailed) are corrected according to the Benjamini-Hochberg (BH) procedure (Benjamini and Hochberg, 1995).

### 5.5 Co-Evolution of interacting protein pairs

We assessed the coevolution of a pair of proteins through a recently released pipeline Miglionico et al. (2024), which first retrieves orthologoues of a given input gene from OMA Altenhoff et al. (2024) and then quantifies the co-presence as the Jaccard similarity of sets of genomes containing orthologues of two proteins.

### 5.6 Cancer evolution

We also retrieved data from cancer clonal evolution from the CancerTracer database (Wang et al., 2020), and assessed the co-evolution, or co-selection, of cancer somatic mutation on interacting gene pairs through the Jaccard similarity of the sets of patients presenting mutations of either gene. The cancer clonal evolution database further divides mutations based on their position in the phylogenetic tree: “trunk”; mutations shared by all samples from a tumor, “branch”; shared by some but not all samples, “private” (which we renamed to “leaf” for consistency), observed in a single sample. We computed the Jaccard index at progressively longer clonal evolution stages starting from the trunk and progressing toward the leaves (i.e. considering trunk only, then trunk and branch together and then the full phylogenetic tree), therefore obtaining three different values for each gene pair.

### 5.7 GPCR signaling axes

We considered GPCRs as a special case study of CCIs. In addition to the ligand-GPCR interactions from the CCIs list, we generated a list of GPCR-G protein couplings by aggregating multiple experimental sources, integrated via GproteinDB (Pándy-Szekeres et al., 2024), as well as predicted couplings from the machine learning predictor PRECOGx (Matic et al., 2022). We only accepted as valid GPCR-G protein pairs those which are predicted to interact by at least 3 different sources. For each ligand-GPCR and GPCR-G protein pair, we computed their co-expression using Pearson correlation across all normal and tumor tissues. Additionally, we assessed their coevolution based on Jaccard similarity, using the orthology-based Jaccard index for normal evolution and the clonal evolution one for cancer. The Kolmogorov-Smirnov *D*-statistic was used to quantify differences between the two interaction types. In particular, the distributions of each metric (co-expression and coevolution) for ligand-GPCR pairs were statistically compared to those for GPCR-G protein pairs using two-sample Kolmogorov-Smirnov tests (scipy, two-tailed, BH-corrected). When considering clonal evolution, we computed the Jaccard index at progressively longer clonal evolution stages starting from the trunk and progressing toward the leaves (i.e. considering trunk only, then trunk and branch together and then the full phylogenetic tree) obtaining three different values for each gene pair.

### 5.8 Hub genes

We considered the total sum of the weights of all links connecting to a gene as that gene’s degree. We then defined hub genes as the top 5% genes by degree in each WGCNA network. For each gene we counted the number of different tissues in which it acts as a hub gene, then we converted the value to a fraction because of the different total number of normal and tumor tissues in our dataset. We measured connectivities of hub genes to gene sets representing biological precesses from MSigDB Hallmark collection (Liberzon et al., 2015), and normalized to 1 the sums of all links having a hub gene as a source or target, ensuring that the results are comparable across different networks regardless of the different values of the WGCNA soft thresholds. When a tissue contains multiple subtissues (e.g. lung adenocarcinoma and lung squamous cell carcinoma), the median value was used.

### 5.9 Rewiring of co-expression modules

We used the signed module eigengene based connectivity measure (KME) (Langfelder and Horvath, 2008) as a fuzzy measure of module membership. By representing each gene by a vector of KME, we measured the tendency of two genes to appear together in a specific tissue/condition combination as the cosine similarity of their KME vectors. This allowed us to consider the co-expression of each gene with all modules eigengenes instead of just assigning a single module to each gene. Only tissues for which both tumor and normal data are available were considered and the cosine similarities were compared using the Wilcoxon signed-rank test, higher cosine similarity values indicate that interacting genes are more likely to appear in the same modules (be co-expressed with the same module eigengenes). In addition, we assessed the overall degree of interaction rewiring within each tissue by calculating the fraction of CCIs that were assigned to the same module in only one condition (tumor same module / normal different modules and vice versa). CCIs assigned to different modules in both conditions were excluded. Sankey plots were generated using the Plotly (v5.19.0) python module to visualize transitions of interactions between modules across conditions. Networkx (v3.2.1) was used to visualize networks of genes involved in *E-M Transition*, CCIs that involve those genes are also included in the network.

### 5.10 Network motifs analysis of the crosstalk network

We aggregated all WGCNA co-expression networks into one by performing Mann-Whitney U between all pairs of proteins or protein complexes that appear in at least one CCI. Specifically, the co-expression of a pair of genes (e.g. a ligand-receptor pair) in a tissue (e.g. heart) was represented by the mean adjacency value of the pair in the WGCNA networks of all subtissues (e.g. left ventricle, atrial appendage). All subtissue networks were normalized so that the maximum degree is 1. When a CCI included multiple genes forming a complex, the co-expression value of that complex with any other complex or gene was represented by the minimum of the co-expression values of all gene pairs involved. All possible pairs of gene/complexes were included, regardless of whether they are actually involved in CCIs or not, but pairs of complexes that share genes were excluded from testing. Test results were summarized by connecting pairs showing significant differences between conditions (*p <* 0.05, two-tailed, BH corrected) with signed links, with +1/-1 values indicating co-expression being greater in tumor/normal tissues.

We studied the topology of the crosstalk network by means of network motifs analysis. The crosstalk network can be thought of as a multiplex network with two layers. The first layer is the co-expression network presented in the previous paragraph, with links representing differences in co-expression across conditions as assessed via Mann-Whitney U testing; links in this layer can exist regardless of the pairs interacting or not. The second layer is the CCI list, with links representing interactions between two genes or complexes, regardless of their co-expression. We started by enumerating all pairs of nodes that are connected in both layers of the network. This corresponds to CCIs showing significant differences in co-expression between normal and tumor. We selected all subgraphs made of two of these interactions. The subgraph representing a pair of interactions includes three nodes if one node is shared by both interactions (e.g. two ligands binding to the same receptor), and four nodes otherwise. We list all possible network motifs these subgraphs can be sorted into based on the connectivity in the co-expression layer of the crosstalk network. We generated all possible isomorphism classes of 3 and 4 nodes and use the *isomorphic* method from the igraph (v0.11.6) python module (Antonov et al., 2023) for detection. To evaluate the enrichment of motifs, we generated a background distribution by means of permutation testing. We considered the CCI list and randomly shuffled the genes/complexes, which corresponds to randomly rewiring the links in the CCI layer of the network, while preserving the degree of each node (e.g. promiscuous receptors will still have many interactions). The co-expression layer of the network is kept fixed. We use 5000 permutations and count the frequency of each network motif in each case. To this end, the occurrences of motifs are normalized by the total number of pairs of interactions that are connected in both layers of the network. Finally z-scores and p-values (BH corrected, two-tailed) are obtained by comparing the frequency of each motif in the actual CCI network with the background distribution of permutations. Additionally, for each clique circuit, we associated the set of biological processes from the MSigDB Hallmark gene set collection (Liberzon et al., 2015) that share common genes with the circuit. Finally, we inspected the DrugBank database (Knox et al., 2010), classifying drugs into antibodies or small molecules. We then identified combinations of two drugs that could potentially target both components of each clique (where one drug targets genes involved in the first CCI and the other targets genes in the second CCI). Only pairs in which both drugs are approved were considered.

### 5.11 Survival analysis

We used Cox-Proportional Hazards (Cox-PH) models to predict the hazard-ratios corresponding to the variations in risk of death between TCGA patient groups. We also generated Kaplan-Meier (KM) plots to visualize survival curves. Cox-PH models are fitted in R (v4.4.2) using the survival package (v3.7) (Therneau, 2020). Survival curves are plotted using the lifelines (v0.30) python (v3.9) module (Davidson-Pilon, 2019). All Cox-PH models are univariate. Survival and censoring information of TCGA patients was retrieved from UCSC Xena, obtaining a total of 29 tumor tissues, and 33 tumor subtypes; no filtering is applied to this dataset. Additionally a pan-cancer analysis is also performed where all tumor subtypes are pooled together. Both in tissue-specific and pan cancer analyses, whenever multiple tumor subtypes correspond to the same tissue (e.g. glioblastoma multiforme and lower grade glioma), a separate baseline hazard function is fit for each tumor subtype. We considered all possible two-CCIs circuits that are significantly more co-expressed in tumor tissues than in normal ones (see previous section). For each pair of interactions, patients are stratified into two groups. The “High Expression” (or “Low Expression”) group contains patients whose expression of each of the genes involved is above (or below) the gene’s median. The two groups are defined independently for each tumor subtype. Patients which don’t fall into one of the groups (some genes are above their median, while some are below) were removed. Cases in which any of the groups contain less than 10 patients were discarded. We selected pairs of interactions with significant differences in survival distribution using the logrank test (*p <* 0.05, BH corrected). Additionally, interactions whose hazard-ratio are compatible with 1 are excluded (error bars correspond to standard error). Next, we identified two-CCs circuits that are better at stratifying patients than individual interactions as those showing decrease in logrank p-value and increase in concordance index. Additionally, the hazard ratio also has to be more distant from the value 1 when using both interactions together than it is for each of the individual interactions. This corresponds to requiring the absolute value of the logarithm of the hazard-ratio of the two-CCIs circuit to be greater than the one of any of the two individual CCIs (e.g. patients stratified based on CXCL11-CXCR3 & EGFR-HBEGF, must obtain a stronger hazard-ratio than both those stratified based on CXCL11 and CXCR3 and those stratified based on EGFR and HBEGF). Finally, we identify processes that are associated with higher or lower survival. Cliques that improve patient stratification are divided into two groups based on the hazard-ratio: good prognosis (HR < 1) and poor prognosis (HR > 1). For each biological process (MSigDB Hallmarks), a contingency table is constructed dividing cliques based on whether they are associated with the process or not, and whether they are linked to good or poor prognosis. Statistical significance of the association is then assessed using Fisher’s exact test (BH corrected, two-tailed p-values).

### 5.12 Immunotherapy response predictions

We assembled a dataset of transcriptomic data from patients that underwent immunotherapy by combining multiple sources, namely data from the TIGER database (Chen et al., 2023), the IMvigor210 cohort (Mariathasan et al., 2018) and TCGA. Microarray data from TIGER is excluded. TCGA treatment data is retrieved from Moiso (2021). All data is converted to log2(FPKM + 0.001) and samples that were collected post-treatment are excluded. Only genes that are detected in all datasets are kept. The data is then split into different tissues, only tissues with at least 20 patients are considered as individual cohorts, resulting in 8 different tissues (skin=474, kidney n=341, bladder n=174, stomach=78, lung n=47, brain n=46, ureter n=24, lymphatic tissue n=22), plus one pancancer cohort obtained by considering all data together regardless of tissue (n=1246, also including 40 patients from other tissues with less than 20 patients each). Each of these cohorts is processed with Combat (Johnson et al., 2007) as implemented in the scanpy (v1.9.6 python module (Wolf et al., 2018) to remove batch-effects. The three different sources of data (TIGER, IMvigor210, TCGA) represent three distinct batches, no other covariates are added. Patient response is converted to a binary variable, complete responders and partial responders are assigned to the “Responsive” (R) category, while all other patients are considered “Non-Responsive” (N).

We employed TabPFN (Hollmann et al., 2025) for predictions due to its speed and effectiveness on small datasets. We evaluated model performance using stratified k-fold cross-validation. For each fold, we trained the model on the remaining folds and obtained predicted probabilities for the held-out fold. By aggregating predictions across all folds, we generated probability estimates for the entire dataset and computed a single value of AUROC and AUPRC. We trained two sets of models, one based on wide sets of genes, one based on individual two-CCIs circuits or pairs of CCIs. In the first case, we considered broad gene sets, namely, all genes, all genes involved in any CCI, all genes appearing in any of the two-CCIs clique circuits, and 11 immunotherapy-related gene signatures obtained from TIGER. When a gene set contains more than 500 features—the maximum input dimensionality allowed by TabPFN—we apply principal component analysis (PCA), retaining components that explain 95% of the variance, to reduce dimensionality. To evaluate model performance, we use a number of cross-validation folds equal to the number of patients in the minority class. This provides the closest approximation to leave-one-out validation while maintaining balanced class distributions across folds. The performance of the different feature sets was compared to the one obtained with all genes using the Wilcoxon signed-rank test (one-tailed p-value). In the case of individual CCIs or two-CCIs circuits, analogously to what we did for the survival analysis, we trained models that combine the genes of two CCIs (e.g. CXCL11, CXCR3, EGFR and HBEGF) and compare the performance with the ones obtained using the genes from each interaction separately (e.g. CXCL11 and CXCR3 or EGFR and HBEGF). Due to the high number of models trained, the number of folds was reduced to 10 in this case. The AUROC of cliques is compared to that of other two-CCIs circuits with the MannWhitney U test (BH corrected, two-tailed). Additionally the comparison was repeated for random pairs of two CCIs (regardless of their co-expression). Finally, we identified two-CCIs circuits that improved immunotherapy predictions. In particular we considered a two-CCIs circuit to perform better than individual CCIs only if using the combination of both interactions increases both the AUROC and the AUPRC with respect to using any of the individual interactions. Out of the predictive two-CCIs circuits, we focused on cliques, assessing the enrichment of biological processes (MSigDB Hallmarks collection) in predictive cliques by permutation testing. First, we counted the occurrences of genes from each biological process in predictive cliques. Next, we replaced the set of predictive cliques with the same number of randomly selected cliques (regardless of their predictive power) and repeat the procedure 5000 times obtaining a background distribution of occurrences of biological processes. Observed values are compared with the background distribution obtaining Z-scores and p-values (BH corrected, two-tailed). We also investigated the predictive power of the expression of the individual *E-M Transition* genes. We considered all genes from *E-M Transition* appearing in at least one clique and obtained AUROCs for their expression (instead of using the prediction probabilities from TabPFN). This allowed us to associate genes with increased (AUROC > 0.5) or decreased (AUROC < 0.5) response rate.

## 7 Acknowledgements

F.R. was supported by the Italian Ministry of University and Research through the Department of Excellence “Faculty of Sciences” of Scuola Normale Superiore. The research leading to these results also received funding from the Italian Association for Cancer Research (AIRC) under *My First AIRC Grant (MFAG) 2020* — ID. 24317 project. F.R. was also supported by the Project granted by *Next Generation EU – National Recovery and Resilience Plan (Piano Nazionale di Ripresa e Resilienza, NRRP) – Mission 4 Component 2 Investment 1.4* – Ministry of University and Research (MUR) Call N. 3277, Project Code ECS_00000017, MUR Directoral Decree n. 1055 (23 June 2022), CUP B83C22003930001, project title “Tuscany Health Ecosystem – THE”, Spoke 8. F.R. also received funding from the ImmunoHub project (“Immunoterapia: cura e prevenzione di malattie infettive e tumorali (Immuno-HUB)”). We gratefully acknowledge the CINECA award, in collaboration with AIRC, for the availability of high-performance computing resources and generous support. We also acknowledge the computational resources of the Center for High-Performance Computing (CHPC) at Scuola Normale Superiore.

## Data and Code Availability

Data from the following resources were used:

- GPCR cancer axes: https://gpcrcanceraxes.bioinfolab.sns.it/
- Cancer Tracer: http://cailab.labshare.cn/cancertracer/
- CellPhoneDB: https://github.com/ventolab/cellphonedb-data
- DrugBank: https://go.drugbank.com/
- IMvigor210: https://github.com/SiYangming/IMvigor210CoreBiologies
- IntAct: https://www.ebi.ac.uk/intact
- Liana: https://github.com/saezlab/liana-py
- MSigDB: https://www.gsea-msigdb.org/gsea/msigdb
- OMA Browser: https://omabrowser.org/oma/home/
- TIGER: http://tiger.canceromics.org
- UCSC Xena Browser: https://xenabrowser.net/

The code used for the analysis was deposited on GitHub at: https://github.com/lorenzoamir/cci_circuits_paper_2025.

## 8 Competing interests

The authors declare no competing interests.

## Notes

### Competing Interest Statement

The authors have declared no competing interest.

### Summary of Updates

Higher quality figures have been included in an updated manuscript version.

